# Generation of a porcine cell line stably expressing pig TMPRSS2 for efficient isolation of viruses from pigs with respiratory diseases

**DOI:** 10.1101/2023.10.14.562371

**Authors:** Yuri L Tanaka, Maya Shofa, Erika P Butlertanaka, Ahmad Massoud Niazi, Takuya Hirai, Hirohisa Mekata, Akatsuki Saito

## Abstract

Pigs are important animals for meat production but can carry several zoonotic diseases, including Japanese encephalitis virus, Nipah virus, and influenza viruses. Several *Orthomyxoviridae* and *Coronavirinae* respiratory viruses require cleavage of envelope proteins to acquire viral infectivity and consequently need a host protease or the addition of exogenous trypsin for efficient propagation. Host TMPRSS2 is a key protease responsible for viral cleavage. Stable expression of human TMPRSS2 in African green monkey-derived Vero cells can enhance the porcine epidemic diarrhea virus. However, considering the narrow host tropism of viruses, a porcine cell line expressing pig TMPRSS2 could be optimal for replicating pig-derived viruses. Herein, we generated and evaluated a pig-derived PK-15 cell line stably expressing pig TMPRSS2. This cell line markedly and specifically enhanced the growth of influenza viruses. Therefore, PK-15 cells expressing pig TMPRSS2 could be a valuable and promising tool for virus isolation, vaccine production, and virological studies.

## INTRODUCTION

Pigs can host various viral infections that can cause significant economic damage to the swine industry [15]-[14]. Pigs can also be a host for several zoonotic viral diseases, including influenza [9], Nipah virus disease [5], and Japanese encephalitis [18]. In particular, pigs can serve as an intermediate host for influenza viruses between birds and humans. To infect cells, human influenza viruses recognize sialic acid (human receptor) linked to galactose (α2-6) while avian influenza viruses recognize sialic acid linked to α2-3 (avian receptor). Previous studies demonstrated that pigs express both human and avian receptors for influenza [23],[16]. Therefore, pigs can be infected with both human and avian influenza A viruses, leading to gene reassortment and the emergence of new strains of influenza that are more likely to spread in humans [21]-[6]. The H1N1 influenza A virus that emerged in 2009 to cause a worldwide pandemic was a porcine-derived influenza virus that spread to humans. Thus, pigs can be an essential host in the spread of infection.

Several respiratory viruses, including *Orthomyxoviridae* and *Coronavirinae*, require cleavage of viral membrane fusion proteins by host proteases to achieve optimal infectivity [7], and trypsin must be added to the culture medium to propagate these viruses. However, since the effect of trypsin is weakened by a culture medium containing high concentrations of fetal bovine serum (FBS), these concentrations must be reduced. This makes subsequent virus isolation and propagation from animal specimens with respiratory symptoms more laborious and complex.

In hosts, TMPRSS2 is key protease that activates respiratory viruses in the respiratory tract [4]. In humans, TMPRSS2 is expressed in a wide range of tissues, including the lung, intestine, and prostate [3]. However, most cell lines do not express TMPRSS2. Previous studies established a Vero cell line with stable expression of human TMPRSS2 [27]. Human metapneumovirus and SARS-CoV-2 can reportedly be isolated and propagated more efficiently in the Vero/TMPRSS2 cell line than in normal Vero cells [27],[25]. Furthermore, the isolation efficiency of porcine epidemic diarrhea virus was also improved by using Vero/TMPRSS2 cells [29]. However, the homology of TMPRSS2 sequences between humans and pigs is low, limiting the use of cells expressing human TMPRSS2 for virus isolation.

Herein, we aimed to establish a pig-derived PK-15 cell line stably expressing pig TMPRSS2 (PK-15/TMPRSS2 cells) to improve the isolation efficiency of respiratory tract-derived viruses. A viral replication assay showed that PK-15/TMPRSS2 cells enhanced the replication of the H1N1 influenza virus >1000-fold than that of normal PK-15 cells. Furthermore, we demonstrated that nafamostat mesylate inhibited viral replication in PK-15/TMPRSS2 cells, suggesting the specific enhancement by pig TMPRSS2. These findings indicate that PK-15/TMPRSS2 cells are a promising tool for isolating and propagating viruses from respiratory or digestive organs.

## MATERIALS AND METHODS

### Plasmids

Plasmids psPAX2-IN/HiBiT and pWPI-Luc2 were kind gifts from Dr. Kenzo Tokunaga [22]. pMD2.G was a gift from Dr. Didier Trono (Cat# 12259; http://n2t.net/addgene:12259; RRID: Addgene_12259).

### Cell culture

Lenti-X 293T (TaKaRa, Cat# Z2180N), PK-15 (Japanese Collection of Research Bioresources Cell Bank (JCRB), Cat# JCRB9040), and PK-15 (*Stat2* k/o) cells [28] were cultured in Dulbecco’s modified Eagle medium (DMEM; Nacalai Tesque, Cat# 08458-16) supplemented with 10% FBS and 1× penicillin–streptomycin (Pe/St; Nacalai Tesque, Cat# 09367-34). VeroE6/TMPRSS2 cells (JCRB, Cat# JCRB1819) [17] were cultured in DMEM - low glucose (Sigma-Aldrich, Cat# D6046-500ML) supplemented with 10% FBS, 20 mM HEPES (Nacalai Tesque, Cat# 17557-94), and 1 mg/mL G-418 (Nacalai Tesque, Cat# 09380-44).

### Viruses

Influenza virus IAV (H1N1) strain A/PR/8/34 (American Type Culture Collection, Manassas, VA, USA, Cat# VR-95) was propagated in specific pathogen-free chicken embryonated eggs. Akabane virus (AKAV; TS-C2 vaccine strain) was purchased from Kyoto Biken Laboratories (Kyoto, Japan) and was propagated in HmLu-1 cells maintained in DMEM with 2% FBS and 1% Pe/St.

The swine influenza virus (SIV, H1N1) used in this study was isolated from nasal swabs of pigs that tested positive for viral RNA (vRNA) by RT-PCR [20]. Nasal swab samples were inoculated into 9–11-day-old embryonated chicken eggs, which were incubated at 37°C for 3 days. Allantoic fluid was harvested from the eggs, and commercial diagnostic influenza A immunochromatography (Fujirebio, Tokyo, Japan) was used to confirm virus isolation. All experiments were performed in a biosafety level 2 facility.

### Determination of the full-length sequence of the SIV (H1N1)

After extraction of nucleic acids from allantoic fluid, RT-PCR was performed using MBTuni-12 and -13 primers complementary to the conserved regions at both ends of the influenza A virus to amplify the full-length sequences of the eight segments of influenza A virus [30]. Purification of RT-PCR products, library preparation, and next-generation sequencing were performed as previously described [19]. Sequences generated by next-generation sequencing were analyzed using CLC Genomics Workbench 11 software (Qiagen). Sequences were then processed to remove primers and low-quality reads and mapped to the reference genomes of SIV strain A/swine/Aichi/101/2018(H1N1) (Acc. Num: MW269569-76) [10]. The SIV (H1N1) sequences determined in this study were submitted to the DNA Data Bank of Japan (DDBJ) under accession no. LC778488-95.

### Generation of a retroviral vector to express TMPRSS2

To generate a retroviral vector expressing pig TMPRSS2, the coding sequence of pig TMPRSS2 was synthesized according to the amino acid sequences deposited in GenBank (Acc. Num: NP_001373060) with codon optimization to pig cells (Integrated DNA Technologies, Inc., Coralville, IA, USA). The synthesized DNA sequence is summarized in the S1 File. Synthesized DNA was cloned into the pDON-5 Neo-vector (TaKaRa, Kusatsu, Japan, Cat# 3657), which was prelinearized with NotI-HF (New England Biolabs [NEB], Ipswich, MA, USA, Cat# R3189L) and BamHI-HF (NEB, Cat# R3136L) using an In-Fusion HD Cloning Kit (TaKaRa, Cat# Z9633N). Plasmids were amplified using NEB 5-alpha F′ Iq competent *Escherichia coli* (NEB, Cat# C2992H) and extracted using the PureYield Plasmid Miniprep System (Promega, Madison, WI, USA, Cat# A1222). The plasmid sequence was verified using a SupreDye v3.1 Cycle Sequencing Kit (M&S TechnoSystems, Osaka, Japan, Cat# 063001) with a Spectrum Compact CE System (Promega).

### Generation of PK-15 cells stably expressing TMPRSS2

Lenti-X 293T cells were cotransfected with pDON-5 Neo-pigTMPRSS2, pGP packaging plasmid (TaKaRa, Cat# 6160), and pMD2.G plasmid with TransIT-293 Transfection Reagent (TaKaRa, Cat# V2700) in Opti-MEM (Thermo Fisher Scientific, Cat# 31985062). The supernatant was filtered 2 days after transfection. Collected retroviral vectors were used to infect normal PK-15 or PK-15 (*Stat2* k/o) cells, which were then cultured in 500 µg/mL G-418 for six days. Single-cell cloning was then performed with a Cell Sorter SH800S (SONY), and the expression of TMPRSS2 in each clone was evaluated via western blotting.

### Rescue of reporter viruses

To rescue an HIV-1–based lentiviral vector, 2.5 × 10^5^ cells Lenti-X 293T cells were cotransfected with 0.4 μg psPAX2-IN/HiBiT, 0.4 μg pWPI-Luc2, and 0.2 μg pMD2.G using three μL TransIT-293 Transfection Reagent in 100 μL Opti-MEM I Reduced Serum Medium. The supernatant was filtered 2 days after transfection. To measure the concentration of Gag p24 protein, the HiBiT value was measured using the Nano Glo HiBiT Lytic Detection System (Promega, Cat# N3040) as described previously [22].

### Preparation of standards for RT-qPCR

Standards for determining copy number of virus stock were prepared using the PrimeScript II High Fidelity One Step RT-PCR Kit (TaKaRa, Cat# R026A) with IAV (H1N1) forward primer (5′-AGACAGCCACAACGGAAAAC-3′) and reverse primer (5′-CTGTTAGGCGGGTGATGAAT-3′), SIV (H1N1) forward primer (5′-TGTTTTTGTGGGGACATCAA-3′) and reverse primer (5′-CCCTTGGGTGTCTGACAAGT-3′), and AKAV forward primer (5′-GGGTGCGTATATGGGTCTTG-3′) and reverse primer (5′-TCTTTGAGTGTAGCGCAGGA-3′). vRNA copy numbers of each virus stock were measured via RT-qPCR assay with the One Step TB Green PrimeScript PLUS RT-PCR Kit (Perfect Real Time) (TaKaRa, Cat# RR096A) as described previously [12]. Primers used were: IAV (H1N1) forward (5′-GGCCCAACCACAACACAAAC-3′) and reverse (5′-AGCCCTCCTTCTCCGTCAGC-3′), SIV (H1N1) forward (5′-TCAAGCCGGAGATAGCAATAAG-3′) and reverse (5′-TTTGTCTCCCGGCTCTACTA-3′), and AKAV forward (5′-GGGTTTCAGAGCCTACAAG-3′) and reverse (5′-GCTACCTCAGGCAACAGATTAG-3′). The PCR protocol was 42°C for 5 min, 95 °C for 10 s, and 40 cycles of 95°C for 5 s, and 60°C for 34 s. Levels of vRNA were normalized to those of vRNA in normal PK-15 cells (ΔΔCt method).

### Virus infection

Normal PK-15 and PK-15/TMPRSS2 #23 cells were seeded into a 96-well plate at 1 × 10^4^ cells per well as previously described [28]; where noted, cells were treated with 100 ng/mL pig IFNβ (Kingfisher Biotech, Cat#RP0011S-025). Cells were incubated for 24 hr. For IAV (H1N1) and AKAV, the culture supernatant was removed, and 100 μL per well virus solution with or without 1μg/mL TPCK-treated trypsin (Sigma-Aldrich, St. Louis, MO, USA, Cat# 4352157) was added per well. After incubation at 37℃ for 2 h, the culture supernatant was removed, and cells were washed once with DMEM + 2%FBS + Pe/St. Subsequently, 150 μL fresh DMEM + 2%FBS + Pe/St was added. Cells were treated with 50, 12.5, 3.13, 0.78, 0.2, 0.05, or 0.01 µM nafamostat mesylate (Selleckchem, Cat# S1386) for 2 hr before infection.

For luciferase-encoding virus, infection was performed as described above. Infected cells were lysed 2 days after infection with a britelite plus (PerkinElmer, Cat#6066769), and the luminescent signal was measured using a GloMax Explorer Multimode Microplate Reader (Promega).

### Quantification of vRNA levels in the culture supernatant

Normal PK-15 or PK-15/TMPRSS2 #23 cells were infected in wells in a 96-well plate (n = 6). After two days, the culture supernatant was collected, and virus RNA levels were measured with RT-qPCR using the One Step TB Green PrimeScript PLUS RT-PCR Kit (Perfect Real Time) as described previously [12]. Briefly, the supernatant was mixed with 2× RNA lysis buffer (2% Triton X-100, 50 mM KCl, 100 mM Tris-HCl [pH 7.4], 40% glycerol, and 0.4 U/μL Recombinant RNase Inhibitor [TaKaRa, Cat# 2313A]) [27]. Primers used were IAV (H1N1) forward (5′-GGCCCAACCACAACACAAAC-3′) and reverse (5′-AGCCCTCCTTCTCCGTCAGC-3′), SIV (H1N1) forward (5′-TCAAGCCGGAGATAGCAATAAG-3′) and reverse (5′-TTTGTCTCCCGGCTCTACTA-3), and AKAV forward (5′-GGGTTTCAGAGCCTACAAG-3′) and reverse (5′-GCTACCTCAGGCAACAGATTAG-3′). The PCR protocol was 42°C for 5 min, 95 °C for 10 s, and 40 cycles of 95°C for 5 s, and 60°C for 34 s. vRNA levels were normalized to those of the vRNA level in normal PK-15 cells, which was used as a control (ΔΔCt method).

### Quantification of vRNA levels in infected cells

Normal PK-15 and PK-15/TMPRSS2 #23 cells were infected as described above. Two days after infection, total RNA was collected using a CellAmp Direct RNA Prep Kit for RT-PCR (Real Time) (TaKaRa, Cat# 3732). GAPDH and vRNA levels were measured with a RT-qPCR assay using the One Step TB Green PrimeScript PLUS RT-PCR Kit (Perfect Real Time). The PCR protocol was 42°C for 5 min, 95 °C for 10 s, and 40 cycles of 95°C for 5 s, and 60°C for 34 s. Primers used were IAV (H1N1) forward (5′-GGCCCAACCACAACACAAAC-3′) and reverse (5′-AGCCCTCCTTCTCCGTCAGC-3′) and *ACTB* forward (5′-TCCCTGGAGAAGAGCTACGA-3′) and reverse (5′-AGCACCGTGTTGGCGTAGAG-3′). vRNA levels were normalized to those of porcine *ACTB*, which was used as an endogenous control (ΔΔCt method).

### Western blotting

To evaluate TMPRSS2 expression, pelleted cells were lysed in 2× Bolt LDS sample buffer (Thermo Fisher Scientific, Cat# B0008) containing 2% β-mercaptoethanol (Bio-Rad, Hercules, CA, USA, Cat# 1610710) and incubated at 70°C for 10 min. TMPRSS2 expression was evaluated using SimpleWestern Abby (ProteinSimple, San Jose, CA, USA) with an anti-Myc tag mouse monoclonal antibody (Cell Signaling Biotechnology, Danvers, MA, USA, Cat# 2276S, ×125) and an Anti-Mouse Detection Module (ProteinSimple, Cat# DM-001). The amount of input protein was visualized using a Total Protein Detection Module (ProteinSimple, Cat# DM-TP01).

### Alignment of TMPRSS2 proteins

The MUSCLE algorithm on MEGA 11 (MEGA Software) was used to align the TMPRSS2 protein sequences from pig (Acc. Num: NP_001373060), human (Acc. Num: NP_001128571.1), cattle (Acc. Num: XP_024835315.1), horse (Acc. Num: XP_014591871.1), dog (Acc. Num: BBD33861.1), cat (Acc. Num: XP_023094478.2), mouse (Acc. Num: AAF97867.1), chimpanzee (Acc. Num: XP_001172064.5), gorilla (Acc. Num: XP_004062887.2), orangutan (Acc. Num: XP_054398540.1), bear (Acc. Num: XP_057162750.1), bat (Acc. Num: XP_037000504.2), rabbit (Acc. Num: XP_051693971.1), elephant (Acc. Num: XP_049730872.1), whale (Acc. Num: XP_007172772.2), sheep (Acc. Num: XP_042093293.1), and camel (Acc. Num: XP_031315750.1).

### Phylogenetic analysis of mammalian TMPRSS2

A phylogenetic tree was constructed using the TMPRSS2 sequences obtained from GenBank. The tree was created using the neighbor-joining method with a general time reversible nucleotide substitution model implemented in a MEGA X program [11]. Bootstrap values >70% (1,000 replicates) are shown.

### Calculation of identity of TMPRSS2 among animal species

The identity of TMPRSS2 sequences among animal species was calculated using MEGA X with a pairwise distance matrix. Analyses were conducted using the Poisson correction model. The rate variation among sites was modeled with a gamma distribution (shape parameter = 5). All ambiguous positions were removed for each sequence pair (pairwise deletion option).

### Statistical analysis

Differences in infectivity between two different conditions (e.g., between normal PK-15 and PK-15/TMPRSS2 #23 cells) were evaluated by unpaired, two-tailed Student’s *t*-test. Differences in infectivity between two different conditions (e.g., between PK-15 (*Stat2* k/o)/TMPRSS2 cells #15, PK-15 (*Stat2* k/o) cells, PK-15/TMPRSS2 #23, and normal PK-15 cells) were evaluated by one-way ANOVA, followed by the Tukey test. p ≤ 0.05 were considered significant. Tests were performed using Prism 9 software v9.1.1 (GraphPad).

## RESULTS

### Generation and screening of PK-15 cells stably expressing pig TMPRSS2

We constructed a phylogenetic tree using the mammalian TMPRSS2 amino acid sequences retrieved from GenBank that shows the genetic difference in the TMPRSS2 sequence between humans and pigs (Figure 1a). The homology rate of TMPRSS2 sequences between humans and pigs is approximately 70%, suggesting that using cells expressing human TMPRSS2 for virus isolated from pig samples is questionable (Figure 1b).

**Figure 1.**
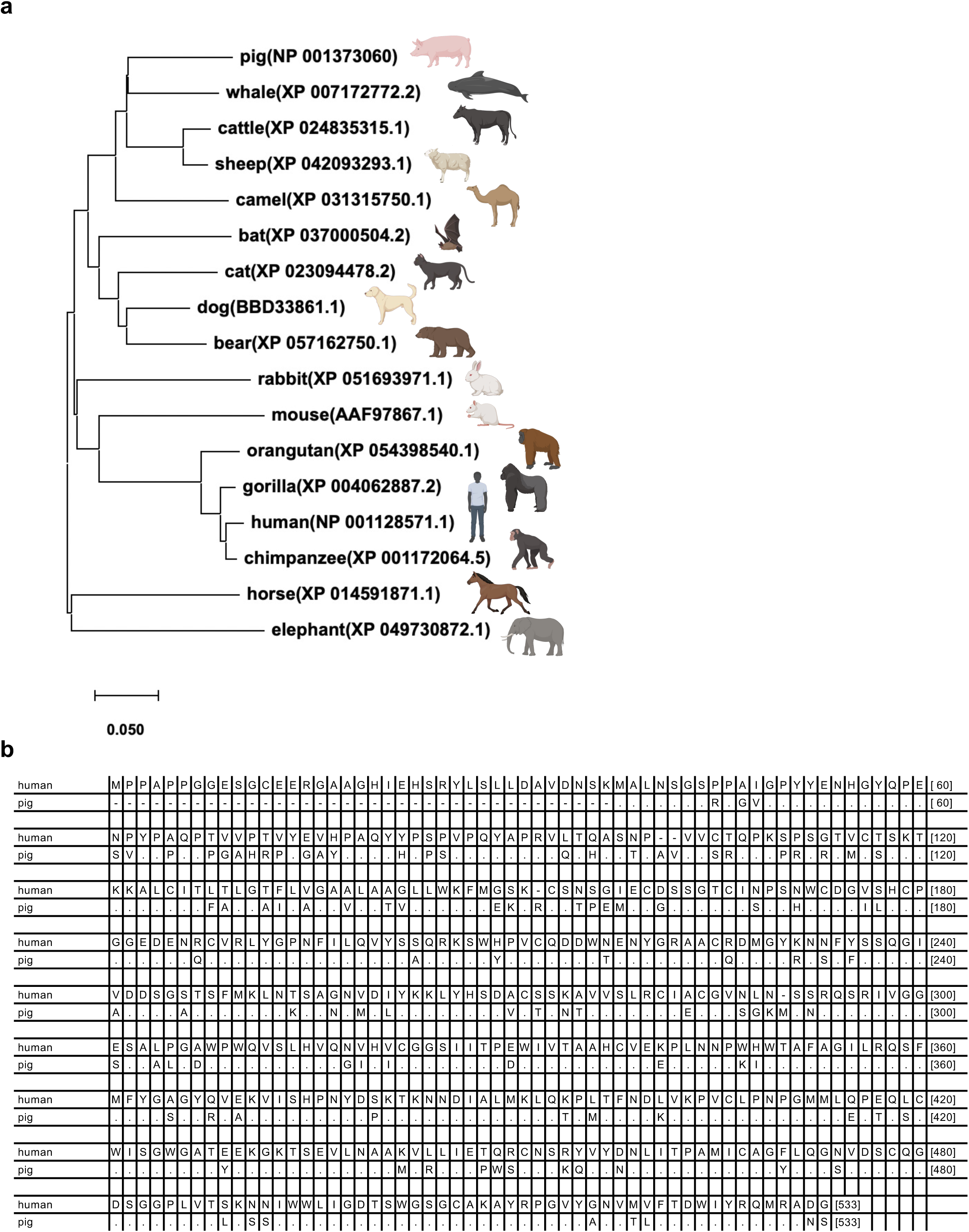
Differences between human and pig TMPRSS2 sequences. (**a**) Phylogenetic tree of TMPRSS2. A phylogenetic tree was constructed using the TMPRSS2 sequences retrieved from GenBank. The tree was constructed using the neighbor-joining method with a general time reversible nucleotide substitution model implemented in a MEGA X program (Kumar et al., 2018). Bootstrap values >70% (1,000 replicates) are shown. (**b**) Alignment of TMPRSS2 amino acid sequences between humans and pigs.

Therefore, we aimed to generate a pig-derived cell line expressing pig TMPRSS2 (pigTMPRSS2) and designed a Myc-tagged pigTMPRSS2 using pig genomic information deposited in GenBank (Acc. Num: NP_001373060) with pig codon optimization. The synthesized DNA was cloned into a retroviral vector that was used to infect normal PK-15 cells, and the transduced cells were selected with neomycin. Single-cell clones were obtained with cell sorting, and western blotting using anti-Myc tag antibody was performed on each cell clone to test the expression levels of pigTMPRSS2 (Figure 2a). We selected PK-15/TMPRSS2 #23 clone because this clone had a higher expression level than that of other clones. Furthermore, we expressed pigTMPRSS2 in PK-15 cells lacking the *Stat2* gene (PK-15 (*Stat2* k/o) cells) [28] to obtain PK-15 (*Stat2* k/o)/TMPRSS2 #15 cells (Figure 2b).

**Figure 2.**
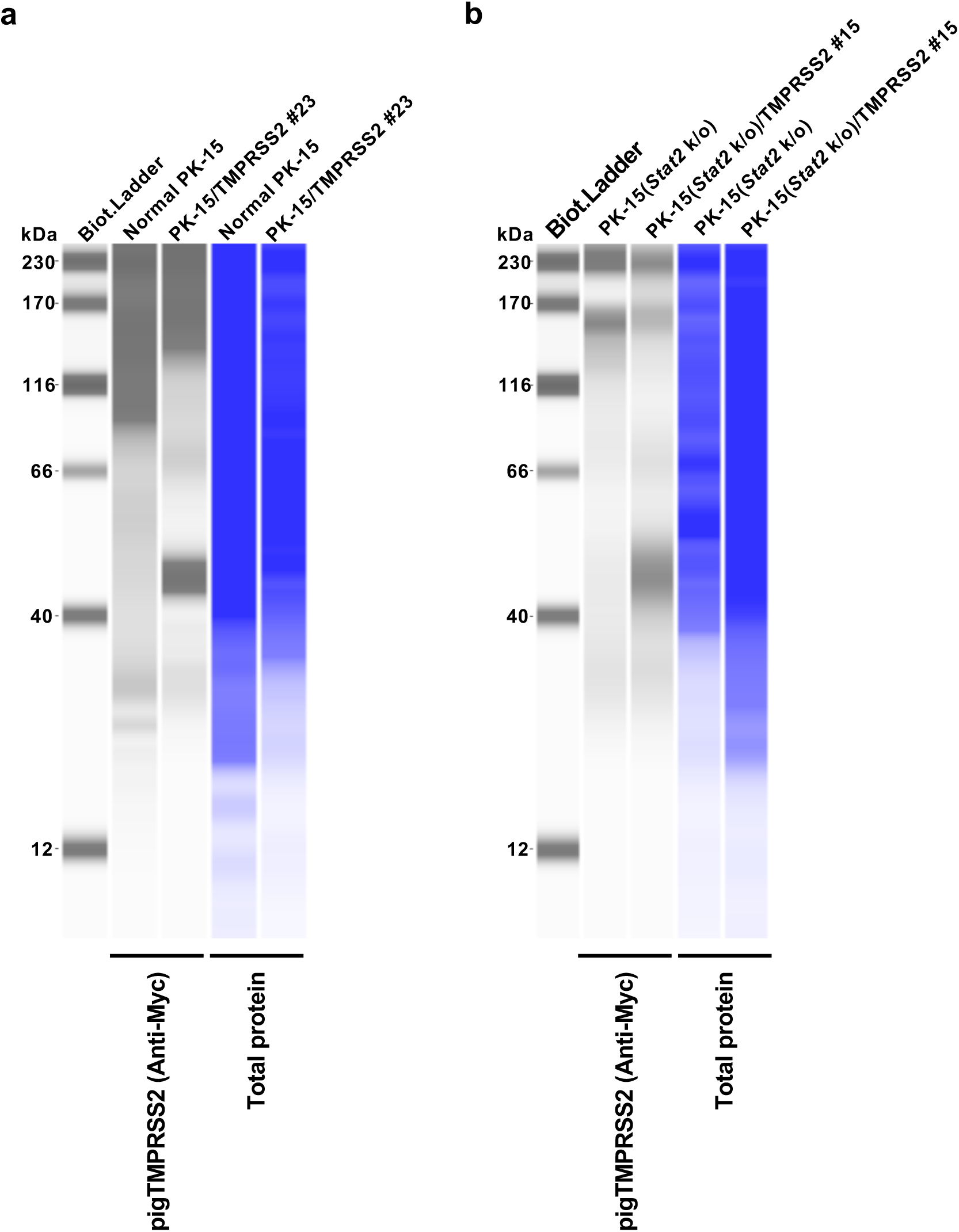
Generation and screening of PK-15 cells stably expressing pig TMPRSS2. (**a**) Expression of the Myc-tagged TMPRSS2 in PK-15 cells as determined with western blotting; the cellular lysate of normal PK-15 cells was used as a negative control (empty). (**b**) Expression of the Myc-tagged TMPRSS2 in PK-15 (*Stat2* k/o) cells as determined with western blotting; the cellular lysate of unmodified PK-15 (*Stat2* k/o) cells was used as a negative control (empty).

### TMPRSS2 expression in PK-15 cells enhances IAV (H1N1) replication

We hypothesized that pigTMPRSS2 expression in PK-15 cells could enhance IAV replication. To address this, we infected normal PK-15 and PK-15 cells expressing pigTMPRSS2 with IAV (H1N1) A/PR/8/34 strain with or without the TPCK-treated trypsin. In normal PK-15 cells, the IAV (H1N1) replication was slightly enhanced by the TPCK-treated trypsin but was significantly enhanced in PK-15/TMPRSS2 #23 cells to approximately 1,000-fold that in the normal PK-15 cells. Furthermore, PK-15/TMPRSS2 #23 cells did not require TPCK-treated trypsin for IAV (H1N1) replication (Figures 3a and 3b). The levels of intracellular vRNA measurements also correlated with those in the supernatant (Figure 3c).

**Figure 3.**
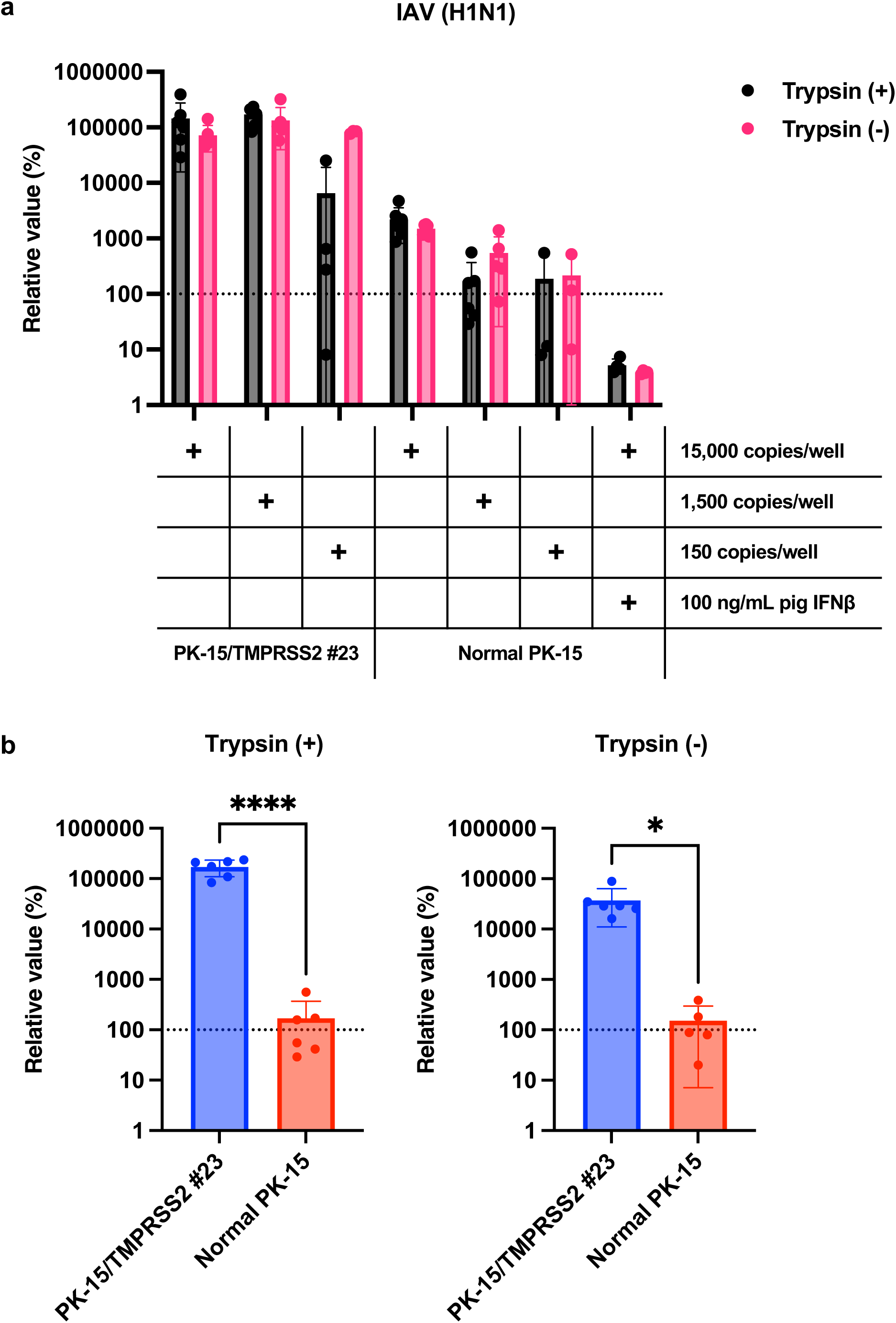
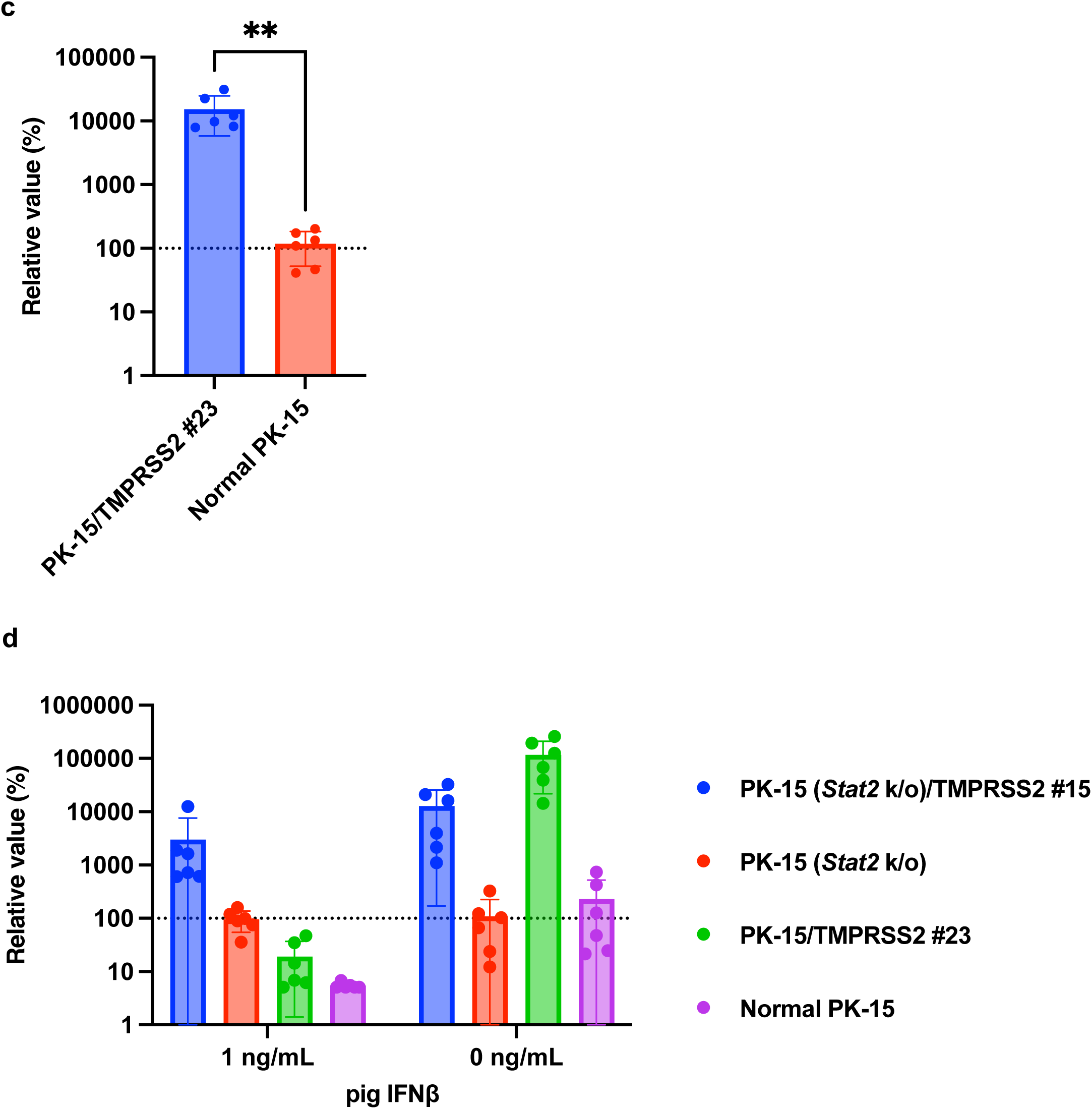
TMPRSS2 expression in PK-15 cells enhances IAV (H1N1) replication. (**a**) PK-15/TMPRSS2 #23 or normal PK-15 cells were infected with 15,000, 1,500, or 150 copies of IAV (H1N1), and vRNA levels in the culture supernatant were measured with RT-qPCR 2 days later. Results are presented as the mean and standard deviation of six measurements from one assay, representing at least three independent experiments. Cells treated with 100 ng/mL pig IFNβ served as negative control. (**b**) The relative value was calculated according to the values in normal PK-15 cells. Results are presented as the mean and standard deviation of six measurements from one assay and represent at least two independent experiments. Differences were examined by a two-tailed, unpaired Student’s *t*-test. \*\*\*\**p* < 0.0001, \**p* < 0.05. (**c**) PK-15/TMPRSS2 #23 or normal PK-15 cells were infected with 15,000 copies of IAV (H1N1), and intracellular vRNA levels were measured with RT-qPCR 2 days later. The relative value was calculated according to the values in normal PK-15 cells. Results are presented as the mean and standard deviation of six measurements from one assay and represent at least two independent experiments. Differences were examined by a two-tailed, unpaired Student’s *t*-test. \*\**p* < 0.01. (**d**) PK-15 (*Stat2* k/o)/TMPRSS2 #15, PK-15 (*Stat2* k/o), PK-15/TMPRSS2 #23, or normal PK-15 cells treated with one ng/mL pig IFNβ or left untreated for 24 hr and then infected with 15,000 copies of IAV (H1N1). vRNA levels in the culture supernatant were measured with RT-qPCR 2 days after infection. The relative value was calculated according to the values in normal PK-15 cells without pig IFNβ treatment. Results are presented as the mean and standard deviation of six measurements from one assay and represent at least two independent experiments.

Interferons (IFNs) exert antiviral effects by binding to the IFN alpha and beta receptor subunit 1 (IFNAR1) on the cellular surface, and STAT2 plays a critical role in this response [24]. Therefore, we tested the usefulness of PK-15 cells lacking *Stat2* for efficient virus isolation from clinical samples in the presence of type I IFNs or IFN-inducible substances such as double-stranded RNA or lipopolysaccharide. Consequently, we observed comparable replication of IAV (H1N1) in normal PK-15 and PK-15 (*Stat2* k/o) cells. We also observed enhanced IAV (H1N1) replication in PK-15/TMPRSS2 #23 and PK-15 (*Stat2* k/o)/TMPRSS2 #15 cells. However, the replication of IAV (H1N1) was significantly blocked when normal PK-15 and PK-15/TMPRSS2 #23 cells were treated with 1 ng/mL pig IFNβ. Conversely, we observed comparable IAV (H1N1) replication in PK-15 (*Stat2* k/o) and PK-15 (*Stat2* k/o)/TMPRSS2 #15 cells compared with that in normal PK- and PK-15 (*Stat2* k/o) cells (Figure 3d). These results suggest that TMPRSS2 expression in PK-15 cells enhances IAV (H1N1) replication. Furthermore, in the presence of IFN or IFN-inducible substances in clinical samples, PK-15 (*Stat2* k/o) cells with stable TMPRSS2 expression may be useful for efficient virus isolation.

### Enhanced SIV (H1N1) replication in PK-15/TMPRSS2 #23 cells

Next, we tested whether the stable expression of pigTMPRSS2 promotes SIV (H1N1) replication. To address this, we infected normal PK-15 and PK-15/TMPRSS2 #23 cells with SIV (H1N1) strain either with or without TPCK-treated trypsin. The SIV (H1N1) replication was significantly enhanced in PK-15/TMPRSS2 #23 cells to approximately 70-fold that in normal PK-15 cells. Furthermore, PK-15/TMPRSS2 #23 cells did not require TPCK-treated trypsin for SIV (H1N1) replication (Figure 4a and 4b). Thus, PK-15/TMPRSS2 #23 cells could be a promising tool for SIV (H1N1) propagation.

**Figure 4.**
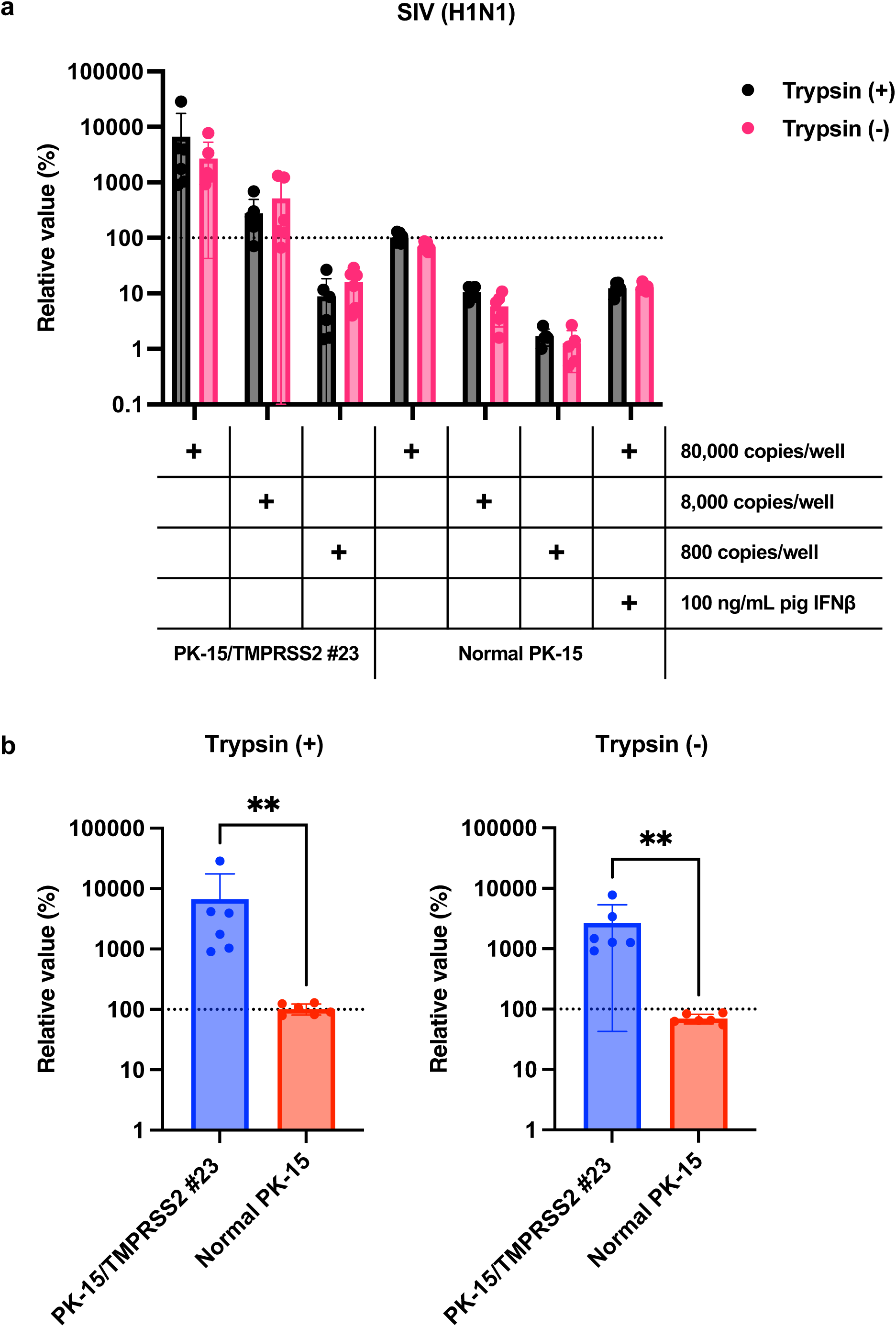
Enhanced SIV (H1N1) replication in PK-15/TMPRSS2 #23 cells. (**a**) PK-15/TMPRSS2 #23 or normal PK-15 cells were infected with 80,000, 8,000, or 800 copies of SIV (H1N1). and vRNA levels in the culture supernatant were measured with RT-qPCR 2 days later Results are presented as the mean and standard deviation of six measurements from one assay, representing at least three independent experiments. Cells treated with 100 ng/mL pig IFNβ served as the negative control. (**b**) The relative value was calculated according to the values in normal PK-15 cells. Results are presented as the mean and standard deviation of six measurements from one assay and represent at least two independent experiments. Differences were examined by a Mann–Whitney *U*-test. \*\**p* < 0.01.

### No enhanced replication of TMPRSS2-indendent viruses in PK-15/TMPRSS2 #23 cells

We then investigated the specificity of enhancement of IAV (H1N1) replication in PK-15/TMPRSS2 #23 cells. We tested whether pigTMPRSS2 expression enhanced AKAV replication since AKAV does not require the TPCK-treated trypsin for viral propagation [2]. AKAV replication between normal PK-15 and PK-15/TMPRSS2 #23 cells did not significantly differ, suggesting that pigTMPRSS2 does not affect AKAV replication (Figure 5a and 5b).

**Figure 5.**
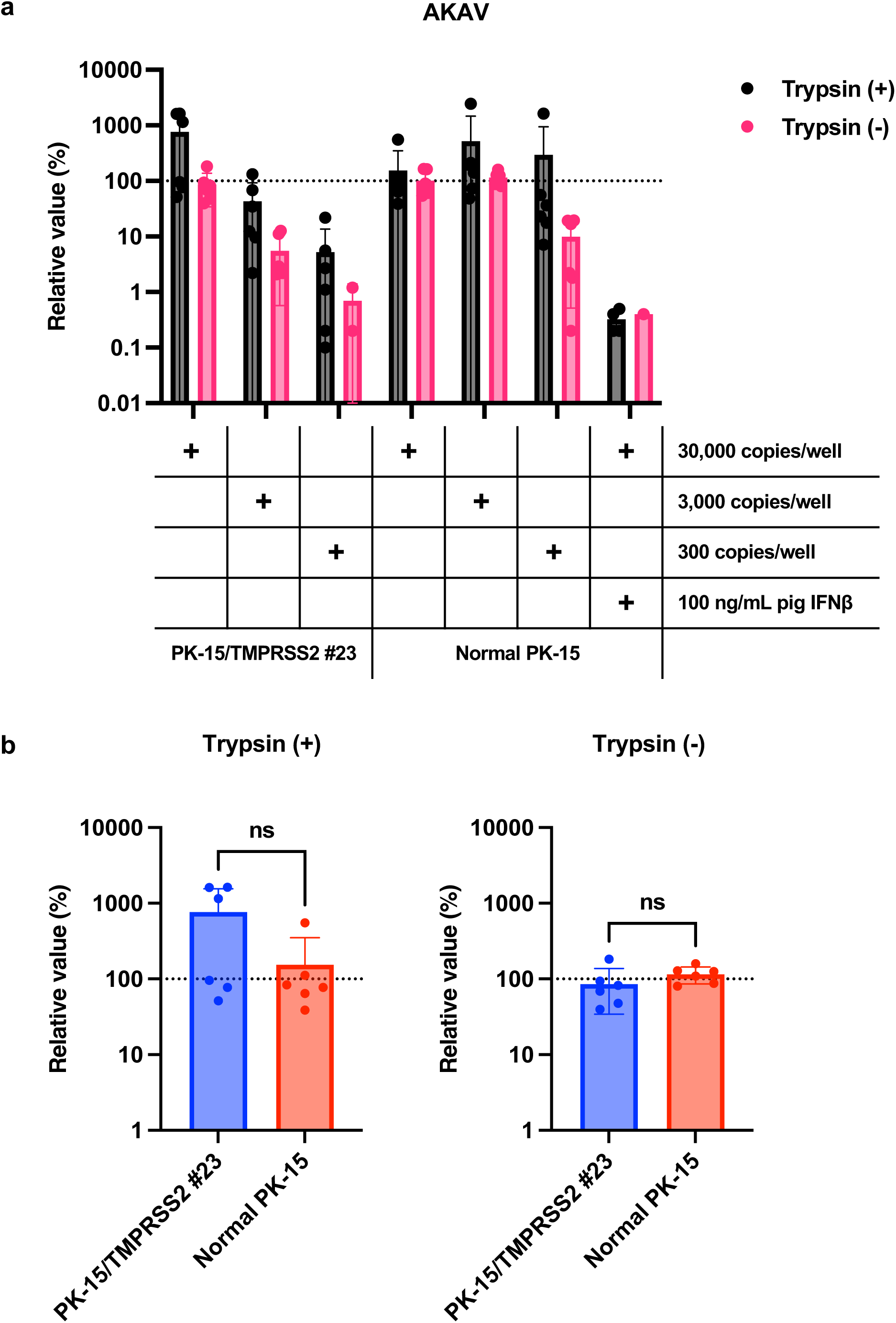
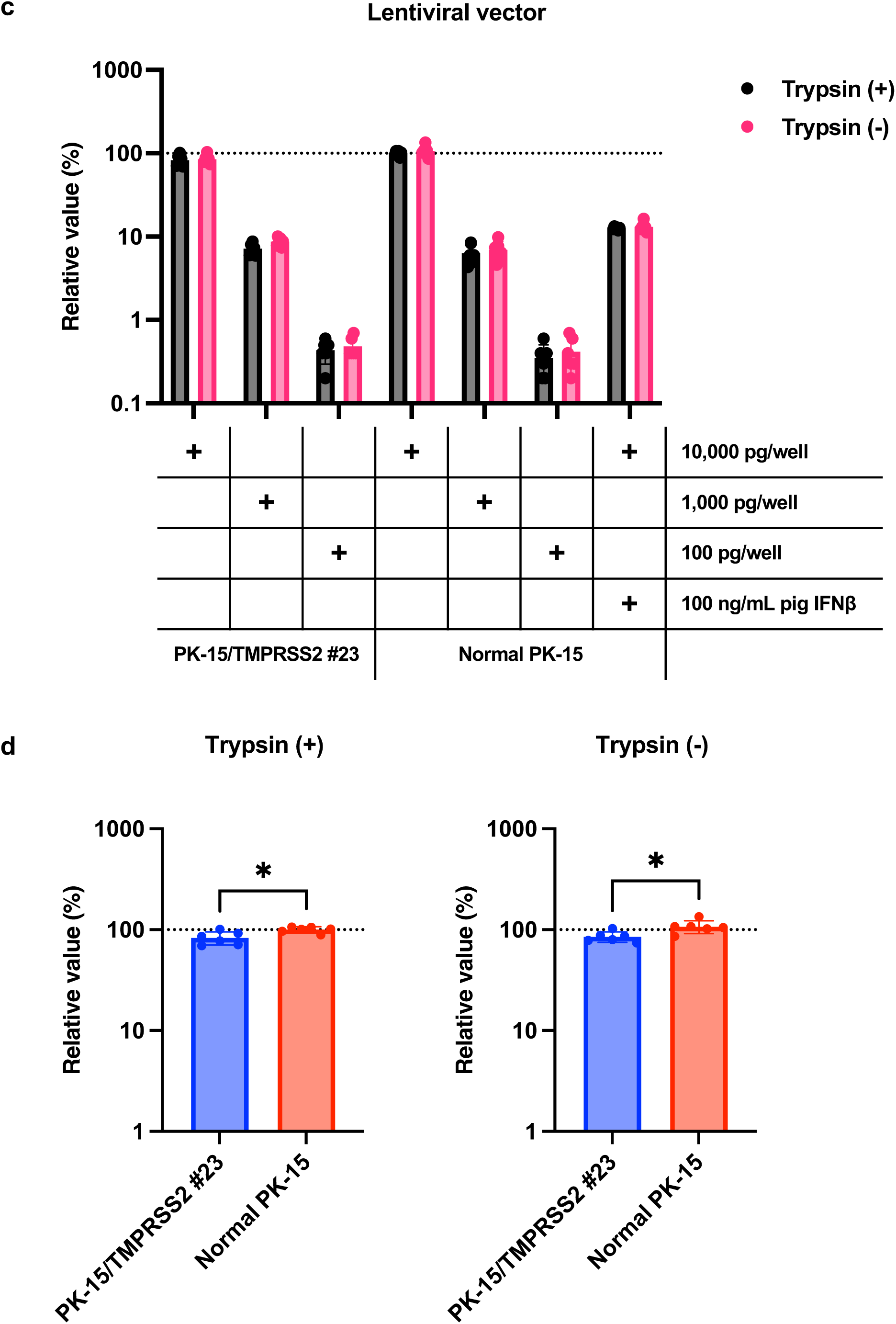
TMPRSS2-indendent virus replication was not enhanced in PK-15/TMPRSS2 #23 cells. (**a**) PK-15/TMPRSS2 #23 or normal PK-15 cells were infected with 30,000, 3,000, or 300 copies of AKAV, and vRNA levels in the culture supernatant were measured with RT-qPCR 2 days later. Results are presented as the mean and standard deviation of six measurements from one assay, representing at least three independent experiments. Cells treated with 100 ng/mL pig IFNβ served as the negative control. (**b**) The relative value was calculated according to the values in normal PK-15 cells. Results are presented as the mean and standard deviation of six measurements from one assay and represent at least two independent experiments. Differences were examined by a two-tailed, unpaired Student’s *t*-test, ns (not significant). (**c**) PK-15/TMPRSS2 #23 or normal PK-15 cells were infected with 10,000, 1,000, or 100 ng (p24) of lentiviral vector, and luminescence was measured 2 days later. Results are presented as the mean and standard deviation of six measurements from one assay, and are representative of at least three independent experiments. Cells treated with 100 ng/mL pig IFNβ served as the negative control. (**d**) The relative value was calculated according to the values in normal PK-15 cells. Results are presented as the mean and standard deviation of six measurements from one assay and represent at least two independent experiments. Differences were examined by a two-tailed, unpaired Student’s *t*-test. \**p* < 0.05.

Finally, we tested whether pigTMPRSS2 enhances the infection of an HIV-1-based lentiviral vector. The lentiviral vector has a VSV-G envelope, and its infection is therefore mediated by endocytosis [1]. No significant difference was present in the lentiviral vector infection efficiency between normal PK-15 and PK-15/TMPRSS2 #23 cells. This suggests that pigTMPRSS2 expression does not affect virus replication and infection of TMPRSS2-independent viruses (Figure 5c and 5d). Therefore, stable pigTMPRSS2 expression in PK-15 cells promotes viral replication of viruses that required the cleavage of envelope proteins by host TMPRSS2.

### IAV (H1N1) replicated more efficiently in PK-15/TMPRSS2 #23 cells than in Vero cells expressing TMPRSS2

Having demonstrated that IAV (H1N1) robustly replicated in PK-15/TMPRSS2 #23 cells, we hypothesized that PK-15/TMPRSS2 #23 cells would be superior to other cell lines for IAV (H1N1) propagation. To investigate this, we compared IAV (H1N1) replication between PK-15/TMPRSS2 #23 and VeroE6/TMPRSS2 cells. IAV (H1N1) replication was significantly higher (approximately 10,000-fold) in PK-15/TMPRSS2 #23 cells than that in VeroE6/TMPRSS2 (Figure 6a). Thus, PK-15/TMPRSS2 #23 cells could be a promising IAV (H1N1) propagation tool.

**Figure 6.**
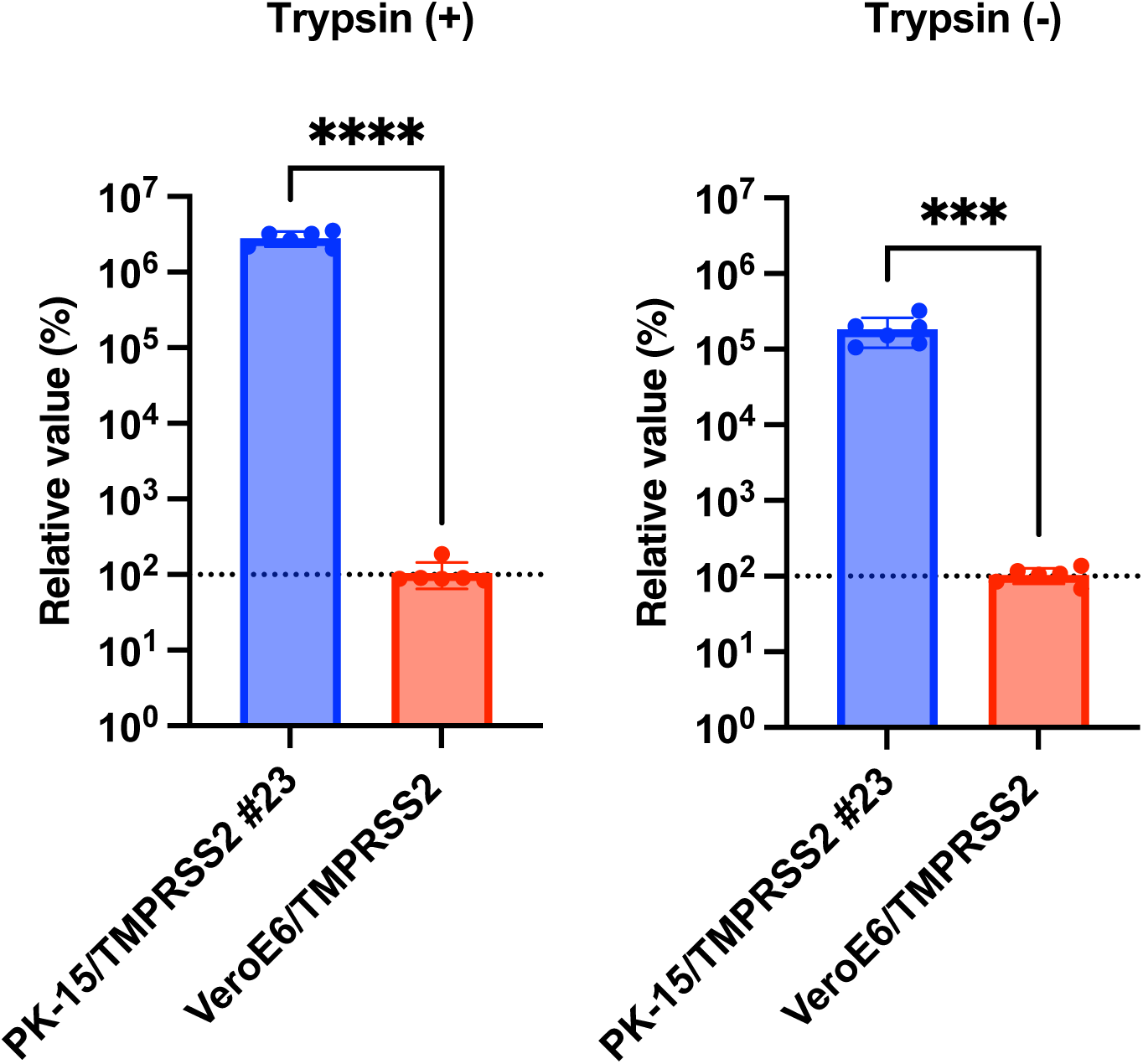
IAV (H1N1) replicated more efficiently in PK-15/TMPRSS2 #23 cells than in Vero cells expressing TMPRSS2. (**a**) PK-15/TMPRSS2 #23 or VeroE6/TMPRSS2 cells were infected with 75,000 copies of IAV(H1N1), and vRNA levels in the culture supernatant were measured by RT-qPCR 2 days later. Results are the mean and standard deviation of six measurements from one assay, representing at least three independent experiments. Cells treated with 100 ng/mL pig IFNβ served as the negative control. The relative value was calculated according to the values in normal PK-15 cells. Results are the mean and standard deviation of six measurements from one assay and represent at least two independent experiments. Differences were examined by a two-tailed, unpaired Student’s *t*-test. \*\*\*\**p* < 0.0001, \*\*\**p* < 0.001.

### PK-15/TMPRSS2 #23 cells can be used for testing of antiviral drugs

Previous studies showed that nafamostat mesylate explicitly inhibits the function of TMPRSS2 [8] and has consequently been used to treat SARS-CoV-2-infected individuals [13]. We therefore tested whether nafamostat mesylate treatment affected the enhanced IAV (H1N1) replication in PK-15/TMPRSS2 #23 cells. Infection experiments performed in the presence of different concentrations of nafamostat mesylate with normal PK-15 and PK-15/TMPRSS2 #23 cells showed that nafamostat mesylate inhibited viral replication in PK-15/TMPRSS2 #23 cells (Figure 7a). The calculated EC_50_ of nafamostat mesylate in PK-15/TMPRSS 2 #23 cells was 0.83651 μM. Conversely, we did not find any effect of nafamostat mesylate in normal PK-15 cells (Figure 7b).

**Figure 7.**
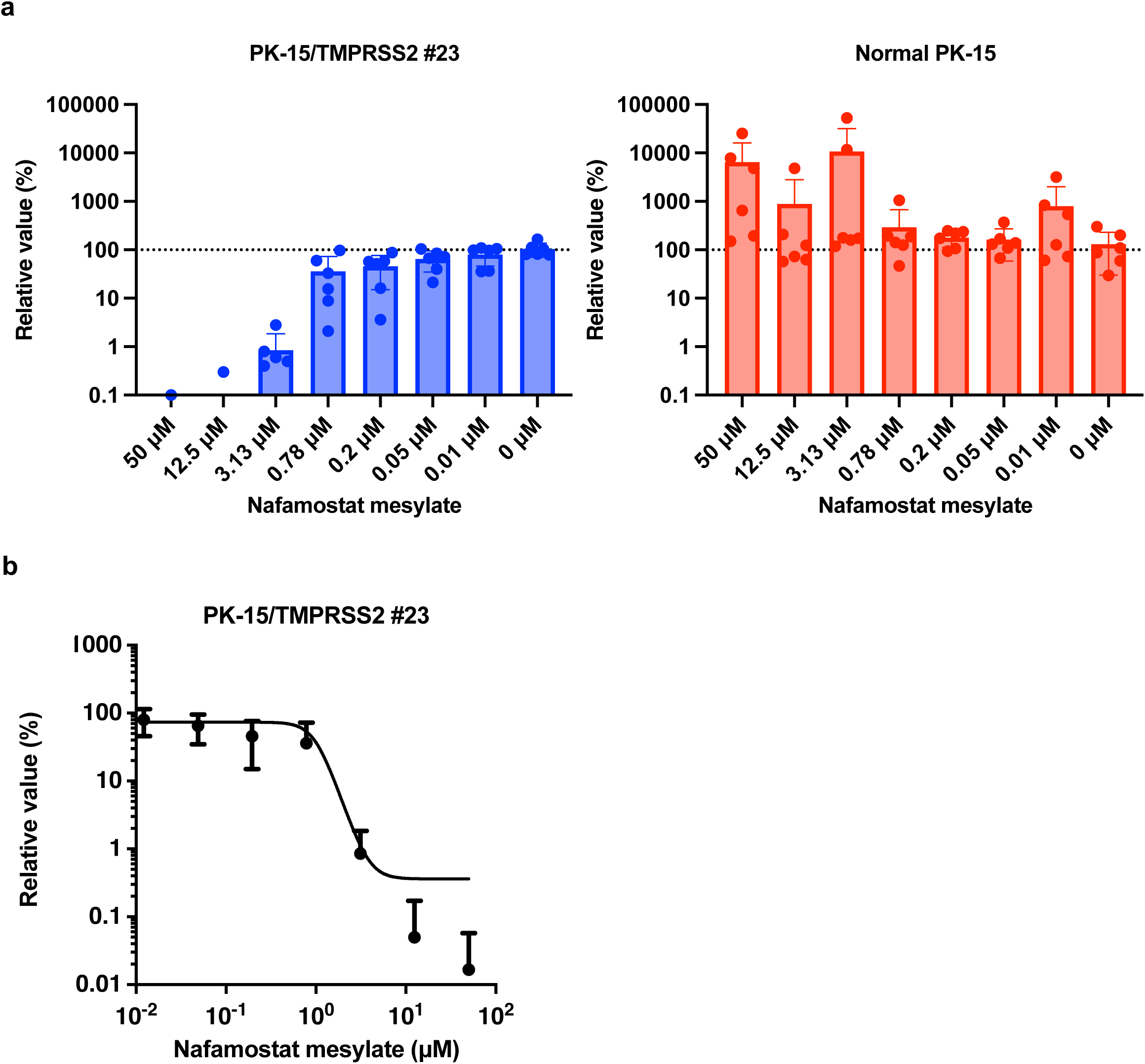
PK-15 cells stably expressing pig TMPRSS2 can be used for testing antiviral drugs. (**a**) PK-15/TMPRSS2 #23 or normal PK-15 cells were infected with 15,000 copies of IAV(H1N1) in the presence of different concentrations of nafamostat mesylate, and vRNA levels in the culture supernatant were measured by RT-qPCR 2 days later. Results are the mean and standard deviation of measurements (n - 6) from one assay and are representative of at least three independent experiments. (**b**) The EC_50_ of nafamostat mesylate against IAV (H1N1) in PK-15/TMPRSS2 #23 cells was calculated using Prism.

Thus, the enhanced IAV (H1N1) replication in PK-15/TMPRSS2 #23 cells was TMPRSS2-dependent, and PK-15/TMPRSS2 #23 cells are therefore a valuable tool for the evaluation of inhibitors against pigTMPRSS2.

## DISCUSSION

Virus isolation from clinical specimens is essential for understanding the growth characteristics and pathogenicity of the virus strain, and the isolated virus can be used as a material to prepare a vaccine antigen. Herein, we generated a porcine PK-15 cell line that stably expressed pigTMPRSS2 to enhance the replication of TMPRSS2-dependent viruses. The PK-15/TMPRSS2 #23 cells significantly enhanced the replication of IAV (H1N1) and SIV (H1N1) even without TPCK-treated trypsin and showed marked inhibition of viral proliferation in the presence of a TMPRSS2 inhibitor. We consider that PK-15/TMPRSS2 #23 cells can improve the viral isolation efficiency from clinical samples from pigs infected with respiratory or digestive tract viruses.

TMPRSS2 plays a significant role in many respiratory viruses, including influenza viruses. Consistent with previous studies [26], the stable expression of TMPRSS2 enhanced the replication of IAV (H1N1) (Figures 3a and 3b) and SIV (H1N1) (Figures 4a and 4b). A characteristic feature of PK-15/TMPRSS2 #23 cells is that we expressed pig TMPRSS2 in pig-derived PK-15 cells. As viruses show optimal replication in their natural hosts, we believe that the PK-15/TMPRSS2 #23 cells can be an essential tool for isolating *Orthomyxoviridae* and *Coronavirinae* viruses from pigs. Furthermore, we successfully generated PK-15 (*Stat2* k/o)/TMPRSS2 #15 cells to isolate viruses in the presence of type I IFNs or IFN-inducible substances. To prove the usefulness of these cells, we will test the isolation efficiency of a variety of pig-derived viruses in future studies.

Interestingly, IAV (H1N1) replicated more efficiently in PK-15/TMPRSS2 #23 cells than in VeroE6/TMPRSS2 cells (Figure 6). The mechanism for this increased IAV (H1N1) replication in PK-15/TMPRSS2 #23 cells remains unclear, although the expression level of TMPRSS2 or difference of activity between human and pig TMPRSS2 may be responsible for this phenotype. Nevertheless, PK-15/TMPRSS2 #23 cells can be a promising tool for isolating and propagating IAV (H1N1) and other viruses.

Our study has some limitations. First, one concern of gene-engineered cells is their specificity. To address this, we demonstrated that PK-15/TMPRSS2 #23 cells specifically enhanced the replications of IAV (H1N1) (Figure 3) and SIV (H1N1) (Figure 4) but not that of AKAV or the lentiviral vector (Figure 5). Furthermore, we showed that nafamostat mesylate canceled the enhanced IAV (H1N1) replication in PK-15/TMPRSS2 #23 cells in a dose-dependent manner (Figure 7). Although these results demonstrated the specific enhancement of TMPRSS2-dependent viruses in PK-15/TMPRSS2 #23 cells, we need to test the applicability of PK-15/TMPRSS2 #23 cells to isolate other viruses. Second, the negative effect of stable TMPRSS2 expression on the isolation or propagation of TMPRSS2-independent viruses should be considered. In this regard, we did not observe any adverse effect of stable TMPRSS2 expression in AKAV replication and infection with lentiviral vectors (Figure 5). Nevertheless, we still need to assess this further in future study.

In conclusion, this study established a pig-derived PK-15 cell line stably expressing pigTMPRSS2. The results clearly demonstrated that PK-15/TMPRSS2 #23 cells could replace exogenous TPCK-treated trypsin for culturing IAV (H1N1), suggesting a promising approach for isolating and propagating TMPRSS2-dependent viruses. We consider that our cells will contribute to the control of viral diseases that are prevalent in pig populations. Furthermore, these cells will be an essential tool to investigate other zoonotic viral diseases including influenza, Nipah virus disease, and Japanese encephalitis.

## ACKNOWLEDGMENTS

pMD2.G was a gift from Dr. Didier Trono. psPAX2-IN/HiBiT and pWPI-Luc2 were kind gifts from Dr. Kenzo Tokunaga. We thank Ms. Tomoko Nishiuchi and Ms. Yuki Shibatani for their support. The authors thank Enago (www.enago.com) for the English language review. Figure 1a was created with BioRender (https://biorender.com/).

This work was supported by grants from the Japan Agency for Medical Research and Development (AMED) Research Program on HIV/AIDS JP23fk0410047, JP23fk0410056, and JP23fk0410058 (to A.S.); AMED Research Program on Emerging and Re-emerging Infectious Diseases JP22fk0108511 and JP22fk0108506 (to A.S.); AMED Japan Program for Infectious Diseases Research and Infrastructure JP22wm0325009 (to A.S.); AMED CRDF Global Grant JP22jk0210039 (to A.S.); from JSPS KAKENHI Grant-in-Aid for Scientific Research (B) 22H02500 (to H.M. and A.S.) and from The Ito Foundation Research Grant R5 KEN77 (to A.S.).

## DATA AVAILABILITY

The datasets of the swine Influenza virus used in the current study are available in the DDBJ (Acc. Num: LC778488-LC778495).

## CONFLICT OF INTEREST

The authors declare no competing interests.

## Notes

### Competing Interest Statement

The authors have declared no competing interest.

