## Supplemental Material 1 for "Generation of a porcine cell line stably expressing pig TMPRSS2 for efficient isolation of viruses from pigs with respiratory diseases"

**Supplemental Materials**

**S1 File. Synthetic DNA for generating plasmids expressing pigTMPRSS2.** The coding sequence of pig TMPRSS2 was synthesized according to the amino acid sequence deposited in GenBank (Acc. Num. NP_001373060.1) with pig cell codon optimization. Start and stop codons are underlined. Letters in lower case indicate the coding sequence of the Myc tag.

ATGgaacagaaactgattagcgaagaagatctgGCACTGAATAGCGGTTCAAGGCCCGGCGTCGGACCATAT
TATGAAAACCACGGCTACCAACCCGAGAGTGTTTATCCACCTCAACCTCCAGGAGCTCACA
GGCCGTACGGAGCCTACCCCGCTCAGTATCATCCGCCCAGCGTTCCACAGTACGCGCCAAG
AGTGCAAACACATGCATCCACACCGGCAGTCGTTGTGTCCCGCCAACCGAAACCCCGCAG
TAGGACGATGTGCAGCTCTAAAACTAAGAAGGCTCTTTGTATTACATTCGCACTTGGAGCA
ATACTCGCAGGAGCGGTCCTTGCGACCGTTCTCCTGTGGAAATTCATGGAGAAGAAAAGA
TGTAGTACGCCTGAGATGGAATGTGGTTCTTCTGGTACTTGCATAAGCCCTTCTCACTGGTG
TGATGGAATCCTTCACTGTCCAGGCGGCGAAGACGAGAATCAGTGCGTCCGGTTGTATGGT
CCTAATTTTATCTTGCAAGTCTACTCCGCCCAACGCAAGTCCTGGTATCCAGTCTGTCAGGA
CGACTGGACTGAGAATTACGGGAGGGCAGCGTGCCAGGATATGGGTTACAGAAACTCTTT
CTTTAGCAGCCAAGGTATCGCCGACGACTCAGGTGCAACTAGTTTTATGAAACTGAATAAA
TCCGCTAATAACATGGATCTGTATAAGAAATTGTACCACAGTGATGTCTGTACTTCCAATAC
GGTGGTCAGTCTTAGGTGTATCGAATGTGGCGTTTCTGGCAAGATGTCTAATCGCCAGTCA
CGGATAGTCGGGGGTAGCAGCGCCGCTCTCGGCGATTGGCCATGGCAAGTTAGCCTCCATG
TGCAGGGGATTCATATATGCGGGGGGTCAATAATTACCCCAGATTGGATCGTGACCGCTGCG
CATTGCGTGGAAGAGCCCCTCAACAACCCTAAAATATGGACGGCGTTCGCAGGTATCTTGA
GGCAATCTTTCATGTTTTATGGCAGTGGATATAGAGTTGCCAAGGTGATCAGTCACCCTAAC
TACGATCCAAAGACTAAAAATAATGATATTGCTCTGATGAAGCTGCAAACGCCTATGACTTT
CAACGATAAAGTTAAACCTGTCTGTCTCCCCAACCCAGGAATGATGCTTGAGCCGACGCAA
TCCTGCTGGATTTCAGGCTGGGGCGCGACATACGAAAAAGGAAAAACCTCAGAGGTTCTC
AACGCCGCTATGGTTCGGCTGATTGAGCCATGGTCCTGTAATAGCAAGCAGGTGTACAATA
ACCTCATAACCCCAGCAATGATATGTGCAGGCTACCTGCAAGGGAGCGTCGATTCCTGCCA
AGGGGATAGCGGAGGCCCCCTTGTCACACTTAAAAGCAGTATATGGTGGCTCATAGGGGAT

ACATCATGGGGATCTGGCTGTGCAAAGGCGTATAGACCGGGCGTGTACGCAAACGTCACA
CTGTTCACAGATTGGATATACAGGCAAATGCGCGCCAATAGTTAA
